# Phospholipid membrane formation templated by coacervate droplets

**DOI:** 10.1101/2021.02.17.431720

**Authors:** Fatma Pir Cakmak, Allyson M. Marianelli, Christine D. Keating

## Abstract

We report formation of coacervate-supported phospholipid membranes by hydrating a dried lipid film in the presence of coacervate droplets. In contrast to traditional giant lipid vesicles formed by gentle hydration in the absence of coacervates, the coacervate-templated membrane vesicles are more uniform in size, shape, and apparent lamellarity. Due to their fully-coacervate model cytoplasm, these simple artificial cells are macromolecularly crowded and can be easily pre-loaded with high concentrations of proteins or nucleic acids. Coacervate-supported membranes were characterized by fluorescence imaging, polarization, fluorescence recovery after photobleaching of labeled lipids, lipid quenching experiments, and solute uptake experiments. Our findings are consistent with the presence of lipid membranes around the coacervates, with many droplets fully coated with what appear to be continuous lipid bilayers. Within the same population, other coacervate droplets are coated with membranes having defects or pores that permit solute entry, and still others are coated with multilayered membranes. These membranes surrounding protein-based coacervate droplets provided protection from a protease added to the external solution. The simplicity of producing artificial cells having a coacervate model cytoplasm surrounded by a model membrane is at the same time interesting as a potential mechanism for prebiotic protocell formation and appealing for biotechnology. We anticipate that such structures could serve as a new type of model system for understanding interactions between intracellular phases and cell- or organelle membranes, which are implicated in a growing number of processes ranging from neurotransmission to signaling.

## Introduction

Biology takes advantage of both lipid self-assembly and biopolymer liquid-liquid phase separation (LLPS) to organize cellular contents, subdividing molecules and reactions via a collection of membrane-bounded and membraneless organelles. Connections between these two general compartmentalization strategies are increasingly recognized to be important.^1, 2^ Intracellular LLPS has been implicated in regulation of transmembrane signaling,^3–5^ autophagy, synaptic signaling,^6^ and cytoskeletal functions.^7, 8^ Intracellular condensates are found in close contact with regions of the plasma membrane and membrane-bounded organelles such as the endoplasmic reticulum.^9, 10^ Functional consequences of LLPS-membrane interactions are just beginning to be understood. For example, LLPS can increase the dwell times of receptor-adaptor complexes at membrane surfaces, effectively providing a noise-filtering mechanism in receptor signaling.^3, 4, 11^

Studies of simpler model systems are a mainstay of research into both lipid membrane and phase-separated droplets. These could aid understanding of how these biophysical organizational motifs may have coevolved and how they interact in living cells. At the same time, inspiration from biological systems can drive advances in synthetic biology, with bioinspired systems designed for new functions. Growing appreciation for membrane-LLPS interactions in vivo has led to some progress in these directions. Submicrometer lipid vesicles (~100 nm diameter) were shown to accumulate at the liquid-liquid interfaces of polyethyleneglycol (PEG)/dextran aqueous two-phase systems^12^ and similarly at the interfaces between complex coacervate droplets and their equilibrium liquids.^13, 14^ Although supported lipid bilayer formation via vesicle fusion on polyelectrolyte multilayer films had been reported,^15, 16^ it was not observed on liquid polyelectrolyte coacervate droplets.^14^ Rather, the vesicles remained intact at these interfaces. The single-chain amphiphile, oleate, when present in solution below its critical micelle concentration, has been shown to self-assemble into multilayered membranes around complex coacervate droplets.^17^ In addition to these examples of assembly at liquid-liquid interfaces, macromolecule-rich phase droplets have been encapsulated within giant phospholipid vesicles as a means to increase the structural complexity and functionality of artificial cells.^18, 19^ This is generally accomplished by first forming vesicles that encapsulate a single-phase macromolecule solution, and then triggering phase separation by a change in temperature, pH, or concentration (the latter via osmotic deflation). As a result, the interior volume is subdivided into two coexisting phases, for example a polyethylene glycol-rich phase and a dextran-rich phase,^20, 21^ or a coacervate phase and a surrounding dilute equilibrium phase.^22–25^ An interesting feature of these two-phase encapsulating vesicle systems is the potential for different interactions between the interior phases and the surrounding membrane, which can lead to adhesion or membrane deformations (vesicle budding, division).^19, 22, 26–29^ A drawback of this approach as a model for intracellular condensate-membrane interactions is the necessity to choose a phase-separating system that can be maintained as a single phase until after lipid vesicles have been assembled, without damage to any of the biomolecules involved due to extremes of temperature or pH. Additionally, if simple lipid hydration or electroformation approaches are used for vesicle production rather than more sophisticated microfluidic methods, the efficiency of encapsulation is quite low (i.e. the vast majority of the total solution remains outside the vesicles), which can be challenging for encapsulation of biomacromolecules that are costly or time-intensive to obtain.^30–35^

Here, we report formation of coacervate-supported phospholipid membranes by hydrating dried lipid films in the presence of pre-existing coacervate droplets. Protein-based and synthetic polymer coacervates showed similar results, in each case producing structures resembling giant lipid vesicles with their entire lumen being a macromolecule-rich coacervate phase. The coacervate-supported membranes were characterized by fluorescence imaging, polarization, fluorescence recovery after photobleaching of labeled lipids, lipid quenching experiments, and membrane permeability experiments that entailed solute uptake and ability to protect against an externally added protease. Our findings are consistent with the presence of lipid bilayers around the coacervates, with some droplets fully coated with what appear to be bilayers while other droplets in the same population have membrane defects or pores that permit solute entry, and others have multilayer membranes. To our knowledge, this is the first example of coacervation templating the formation of phospholipid membranes.

## Results and Discussion

### Coacervate-templated membrane formation

We compared giant vesicle formation in the buffer versus in the presence of coacervate solution as hydrating solution (Figure 1A, B). In our initial experiments, we took advantage of the upper critical solution temperature (UCST) of protamine sulfate coacervates to generate a single-phase solution, which we then used as the hydrating liquid for lipid vesicle formation via the gentle hydration method.^36, 37^ When 20 mM protamine sulfate in 5 mM HEPES and 1 mM MgCl_2_ at pH 7.4 and 40 °C was added to a dried film of (37.9 mol% DOPC, 30 mol% DOPE, 30 mol% DOPS, 2 mol% DOPE-PEG 2kDa, 0.1 mol% DOPE-Rh), the initially clear solution instantly became cloudy, suggesting phase separation despite the elevated temperature (T > UCST). Subsequent observation of these samples via optical microscopy at T<UCST showed coacervate droplets surrounded by a thin layer of fluorescently labeled lipid, lowering the critical temperature for phase separation and facilitating lipid assembly at the coacervate/dilute phase interface. Since coacervation had appeared to be immediate, we next tested whether pre-existing coacervates added to a dried lipid film would become coated with lipid. Figure 1C shows the results of experiments in which protamine sulfate coacervates in 1 mM MgCl_2_, 5 mM HEPES pH 7.4 buffer were added to a dried lipid film (37.9 mol% DOPC, 30 mol% DOPE, 30 mol% DOPS, 2 mol% DOPE-PEG 2kDa, 0.1 mol% DOPE-Rh) and incubated at 40 °C for 2 days. In these samples, fluorescently labeled lipid was found almost exclusively at the protamine sulfate coacervate droplet-supernatant phase interface (Figure 1C, left panels). For comparison, vesicles formed by gentle hydration of an identical lipid film within the same buffers that were used to form coacervates are shown in Figure 1C, right panels. The lipid structures formed in the presence of coacervates are much more uniform in overall size, uniformity of lipid structure thickness, and shape as compared to those hydrated without coacervates, a result of the coacervate droplets acting to template lipid assembly. When samples were observed in real-time during hydration, we were able to capture a rapid process of membrane formation around a coacervate droplet, in which an existing patch of membrane at the coacervate/dilute phase interface grew to ultimately encapsulate the entire droplet (Figure 1D).

**Figure 1:**
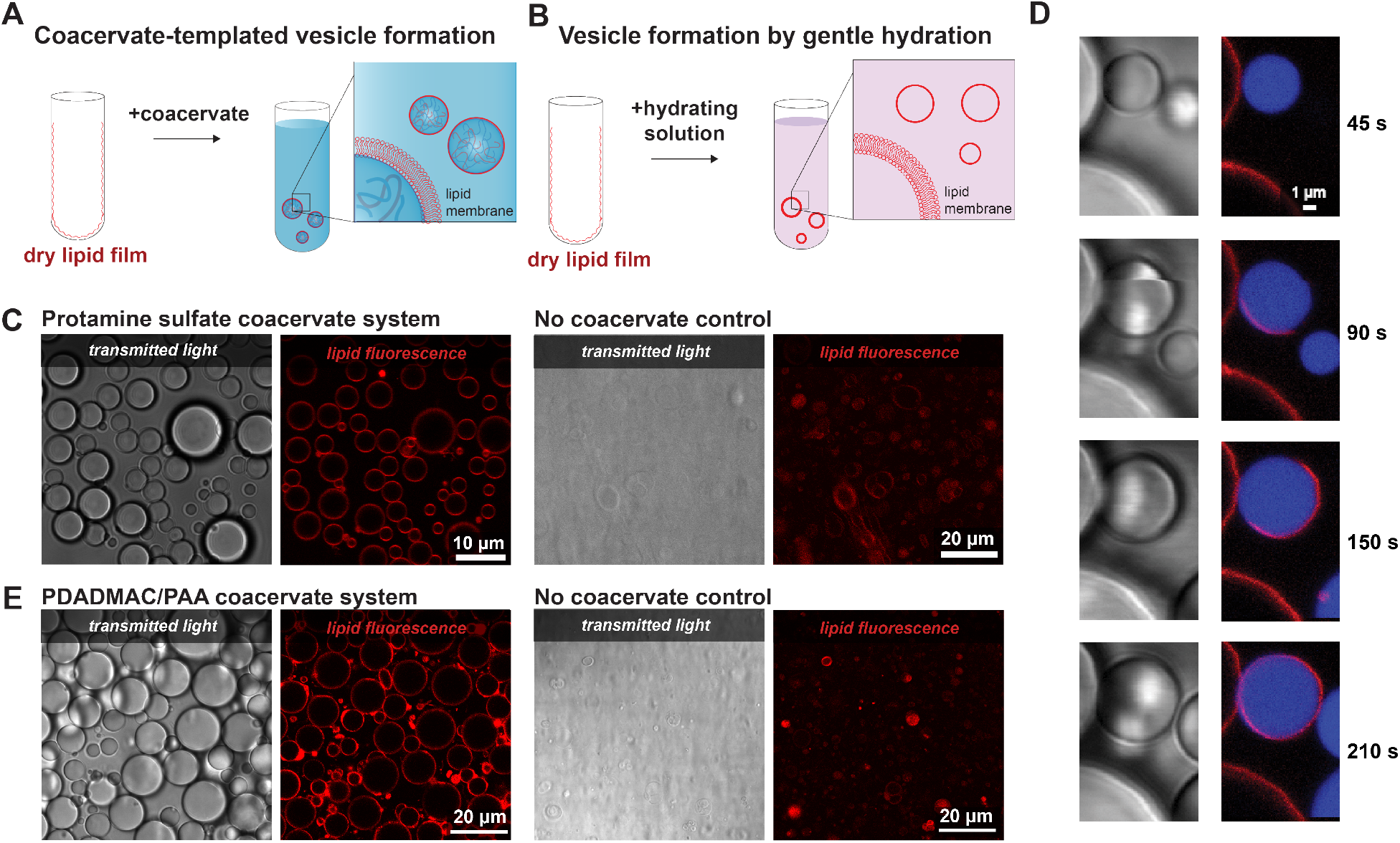
Coacervate templated membrane formation. (A) Schematic illustrating coacervate-templated vesicle formation via gentle hydration in the presence of coacervate droplets. (B) Schematic illustrating traditional vesicle formation via gentle hydration with no coacervates present. (C) Microscope images of coacervate-templated vesicles generated in protamine sulfate coacervate system (left panels) as compared with a no-coacervate control sample (right panels). Lipid composition was 37.9 mol% DOPC, 30 mol% DOPE, 30 mol% DOPS, 2 mol% DOPE-PEG 2kDa, 0.1 mol% DOPE-Rh in 1 mM MgCl_2_, 5 mM HEPES pH 7.4 buffer. (D) Transmitted light (left) and fluorescence (right) images of the membrane formation around protamine sulfate coacervate droplets over time. DOPE-Rh fluorescence and labeled RNA U 15 are false-colored (red and blue, respectively) to aid in visualization. (E) Microscope images of coacervate-templated vesicles generated in PDADMAC/PAA coacervate system (left panels) as compared with a no-coacervate control sample (right panels). Lipid composition was 25 mol. % POPS, 50 mol. % POPC and 22.9 mol. % POPE, 2 mol.% DOPE-PEG 2kDa and 0.1 mol.% DOPE-Rh in 100 mM NaCl, 10 mM Tris pH 7.4 buffer.

The formation of lipid membranes around coacervate droplets was not limited to the specific choice of coacervate-forming polymers or lipid composition: repeating the experiment with synthetic polymer coacervates formed from poly(diallyldimethylammonium chloride) (PDADMAC)/ polyacrylic acid (PAA) with slightly different lipid composition (25 mol. % POPS, 50 mol. % POPC and 22.9 mol. % POPE, 2 mol.% DOPE-PEG 2kDa and 0.1 mol.% DOPE-Rh) in 100 mM NaCl, 10 mM Tris pH 7.4 buffer., the outcome was essentially the same (Figure 1E). Comparing the samples prepared with and without coacervates, the overall yield of apparent giant lipid vesicles increased by roughly tenfold (9-fold for protamine sulfate and 12-fold for PDADMAC/PAA; see Figure S1 and Supporting Information for details). Over 95% of vesicles contained coacervate phase, strongly suggesting that the coacervate droplets template lipid membrane formation (Table S1). From the optical microscopy images of Figure 1, it is apparent that the coacervates are coated in some form of lipid membrane. In the next sections, we perform experiments to better understand the nature of these interfacial lipid assemblies.

### Membrane properties

When fluorescently-tagged lipids are organized into the membranes of traditional giant unilamellar vesicle, differences in fluorophore orientation as a function of location around the vesicle leads to polarized fluorescence that is indicative of the degree of orientation and is not observed for randomly-oriented dyes.^38–40^ Higher polarization correlates to higher molecular packing and alignment, while lower polarization correlates to more fluid, liquid-like membranes.^38, 39, 41^ The lipid-coated coacervate droplets showed strong polarization in the emission of rhodamine-labeled DOPE, with intensity in the y-direction more than double that in the x-direction (Figure 2 A and C), indicating the alignment of lipid molecules at the interface, which is consistent with the presence of one or more well-organized bilayer membranes. In contrast, in previous work where small (~100 nm) unilamellar lipid vesicles assembled at the interfaces of poly(ethylene glycol)/dextran aqueous two-phase systems (ATPS)^12, 42^ or coacervate droplets^13^, no polarization was observed.

**Figure 2:**
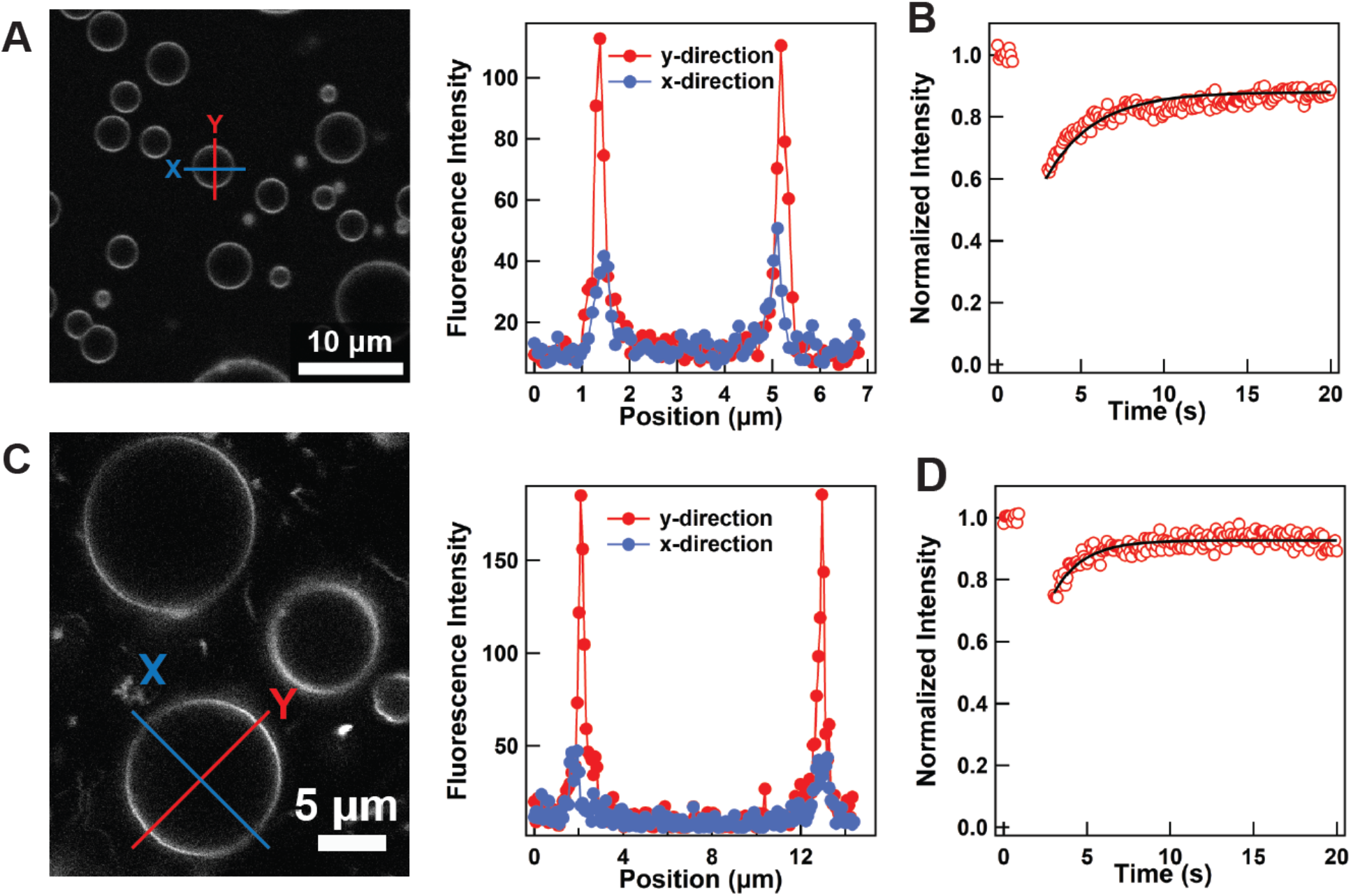
Polarized fluorescence and rapid diffusion of coacervate-templated membranes. (A) and (C) fluorescence confocal image and intensity plots showing increased fluorescence intensity in the y-direction compared to that in the x-direction, confirming fluorescence polarization of the membrane around protamine sulfate and PDADMAC/PAA, respectively. (B) and (D) FRAP recovery data (red circles) for membrane fit (black trace) for protamine sulfate and PDADMAC/PAA, respectively. Lipid composition of 37.9 mol% DOPC, 30 mol% DOPE, 30 mol% DOPS, 2 mol% DOPE-PEG 2kDa, 0.1 mol% DOPE-Rh was used for protamine sulfate in 1 mM MgCl_2_, 5 mM HEPES pH7.4 buffer and lipid composition of 25 mol. % POPS, 50 mol. % POPC and 22.9 mol. % POPE, 2 mol.% DOPE-PEG 2kDa and 0.1 mol.% DOPE-Rh was used for PDADMAC/PAA in 100 mM NaCl, 10 mM Tris pH 7.4 buffer.

Individual lipid molecules within a fluid phase lipid bilayer diffuse laterally,^43, 44^ therefore if one or more bilayers are present around the coacervate droplets, we would expect to see fluorescence recover after photobleaching a portion of the labeled interfacial lipid. Fluorescence recovery after photobleaching (FRAP) was recorded for both lipid membranes around both protamine sulfate and PDADMAC/PAA coacervates (Figure 2 B and D). Almost full recovery was observed for both systems, indicating that lipids in the membrane were mobile.^45–47^ This observation, together with the polarized fluorescence, rules out the possibility that interfacial lipid exists as subresolution liposomes^13^ or lipid aggregates, and is consistent with lipid bilayer membranes. The calculated apparent diffusion (D_app_) constants were 0.10 ± 0.045 μm^2^ and 0.30 ± 0.18 μm^2^ for protamine sulfate and PDADMAC/PAA, respectively. These values are within the range reported values for cell membranes (0.8 ± 0.06 μm^2^) and nuclear membranes (0.41 ± 0.1 μm^2^).^48, 49^ More rapid diffusion has been reported for supported lipid bilayers (1.9 ± 0.3 μm^2^) and GVs (3.7 ± 0.5 μm^2^) formed with DOPC phospholipid with DOPE-Rh label.^50^ These values decreased when polymer supported systems were used.^15^ We hypothesize that the difference for Dapp between our two systems is a result of the different interactions between lipids and polymer/protein system used.

To further evaluate the possibility of lipid bilayer formation around the coacervate droplets, we next performed quenching experiments for fluorescently labeled lipid in outer leaflet. The PDADMA/PAA coacervate templated GVs system was used, with fluorescent NBD labelled lipid (NBD-PE) introduced to the membrane-forming POPC/POPS/POPE lipid mixture. When dithionite, which is unable to permeate intact lipid membranes, is introduced to the external solution, it irreversibly quenches the outer leaflet NBD dye, but cannot reach NBD in the inner leaflet of the membrane.^51^ Taking advantage of slow transverse (flip-flop) diffusion rates,^52^ this experiment can evaluate membrane lamellarity.^53^ When dithionite was added to the lipid-coated coacervates, we observed a decrease of fluorescence intensity of roughly 1/2 (quenched 0.49 ± 0.12 and reference 1 ± 0.18) (Figure 3), consistent with the presence of a lipid bilayer membrane around the coacervate droplets. For quenching calculations, coacervate templated membranes with likely multiple layers indicated by much higher fluorescence intensities were excluded both in reference and quenched systems (see Figure S2 for an example).

**Figure 3:**
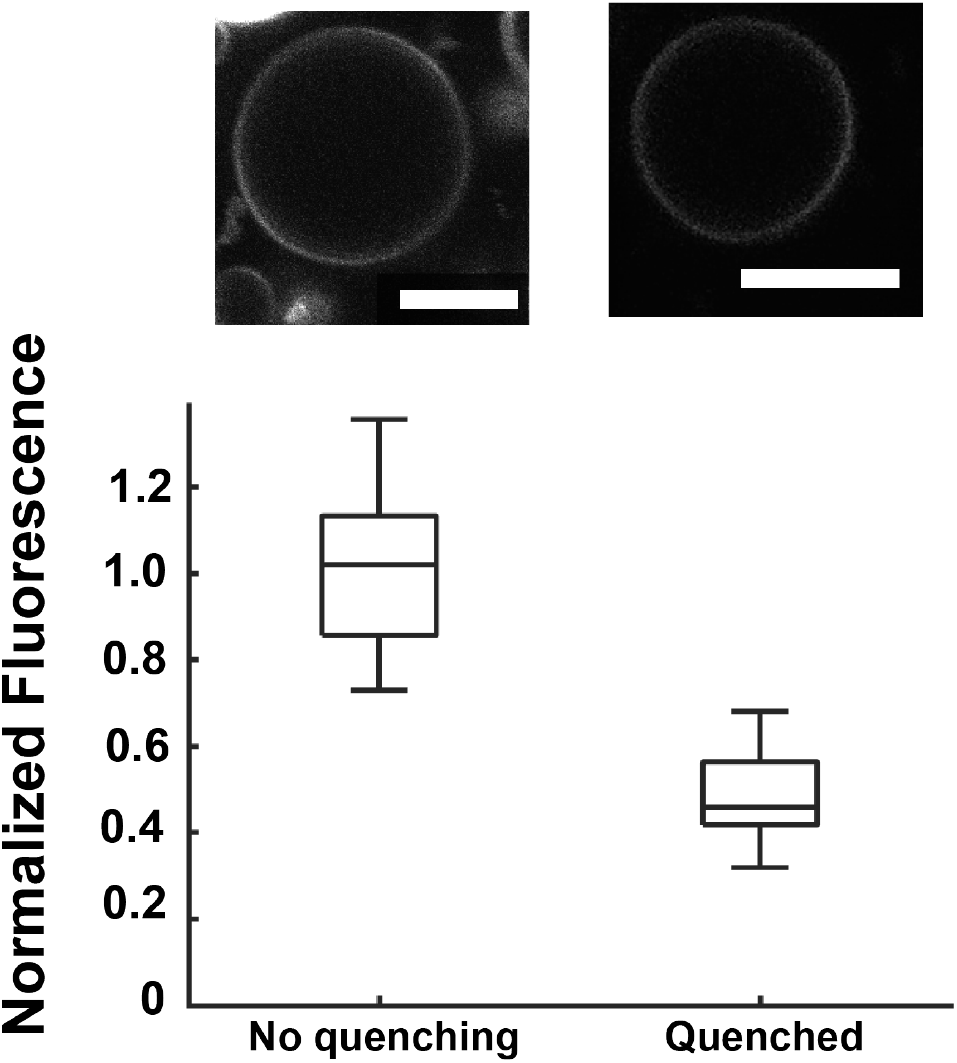
Fluorescent intensities after and before quenching experiment of NBD-PE in the outer leaflet. Representative unedited fluorescence images of coacervate templated membrane after ~1-hour quenching (left) and before quenching (right) were shown. PDADMAC/PAA coacervate system with 25 mol. % POPS, 50 mol. % POPC and 22.9 mol. % POPE, 2 mol.% DOPE-PEG 2kDa and 0.1 mol.% DOPE-Rh lipid composition in 100 mM NaCl, 10 mM Tris pH 7.4 buffer. Scale bars 5 μm. Center lines show the medians; box limits indicate the 25th and 75th percentiles; whiskers extend to the most extreme data points. Error bars show the standard deviation of ~ 21 droplets obtained.

### Membrane barrier

An important function of lipid membranes is to provide a barrier to solute entry/egress.^54^ We investigated the permeability of our coacervate-supported membranes to both the small molecule fluorescein isothiocyanate (FITC) (MW 390 Da) with a negative charge^55^ and the fluorescently-labeled oligonucleotide U15-Alexa 647 (U15) (MW 4860 Da) (Figure 4). Coacervate systems can concentrate a wide range of solutes including small molecules, nucleic acids and other biomolecules.^13, 56^ When fluorescent solutes are added to the exterior solution of uncoated coacervates, they rapidly accumulate into the coacervate droplets (Figure S3). When the same experiment is performed for membrane-coated coacervates, some coacervates accumulate labeled solute, while others remain dark. These data indicate that some coacervate-supported membranes are more fully intact than others, with the more permeable membranes likely having pores and/or patches of incomplete or imperfectly organized membrane (Figure S4). We note that some cationic peptides, particularly oligoarginines, can function as cell-penetrating peptides,^57, 58^ and that the protamine has a high arginine content.^36^ The variability in solute uptake between droplets despite the uniform protamine content of their coacervate interiors points to explanations other than potential cell-penetrating properties of the cationic coacervate components. For the datasets shown in Figure 4, approximately 52% and 57% of the membrane-coated protamine sulfate and PDADMAC/PAA coacervates, respectively, excluded FITC. This suggests that for this slight majority of the population, lipid membrane formation was complete, without pores or defects large enough to allow this small molecule to pass (Figure 4B-D). When an RNA oligomer, a fluorescently labeled 15-mer of uridylic acid (U15), was added to the external solution, it was excluded from approximately 69% and 62% of the membrane coated protamine sulfate and PDADMAC/PAA coacervates, respectively (Figure 4A-C). The greater population of membrane-coated droplets exhibiting good barrier function for the U15 as compared to FITC may be due to the greater molecular weight and higher charge of the U15 RNA, which would necessitate a larger disruption in bilayer structure for permeability.

**Figure 4:**
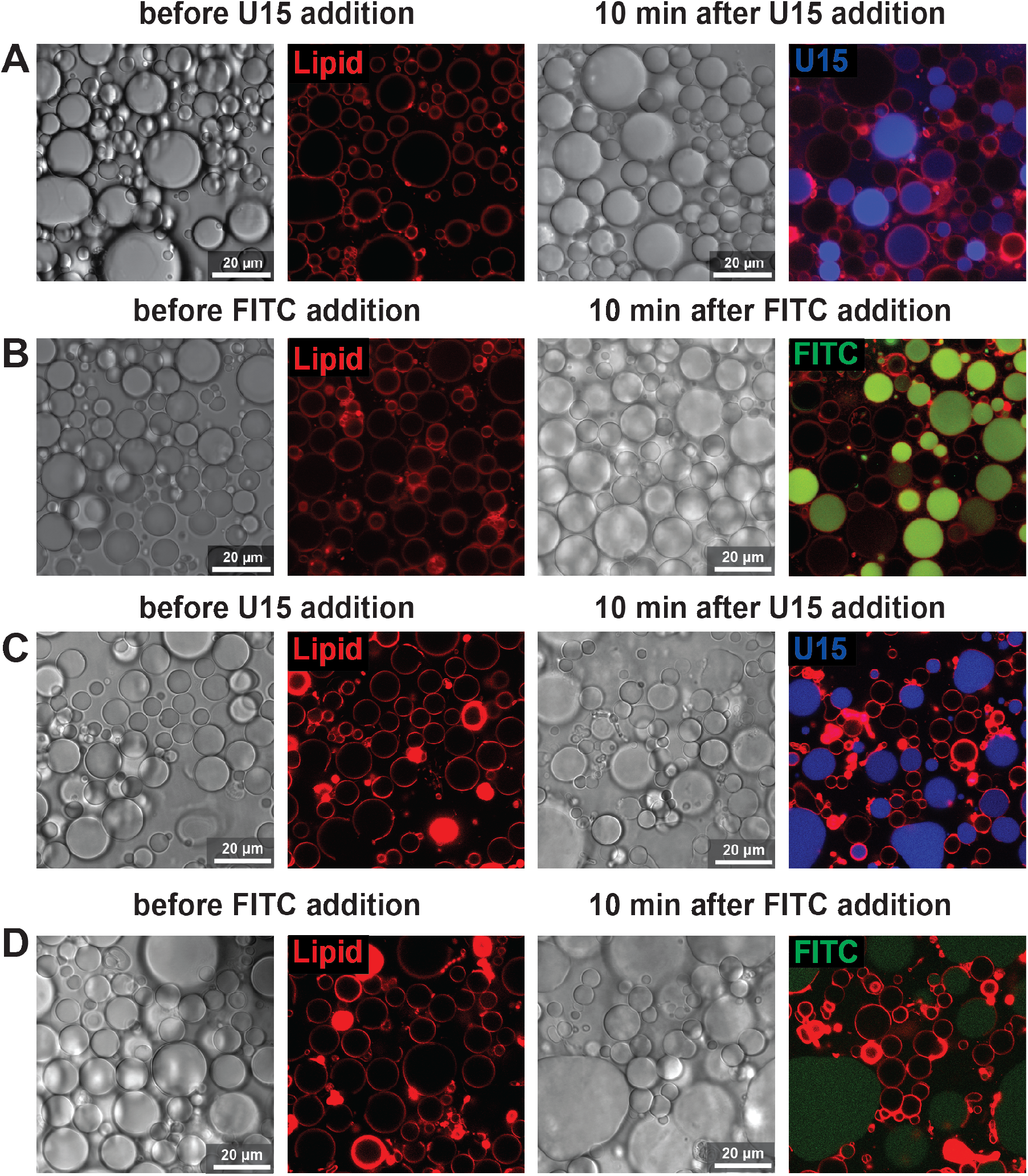
Permeability of coacervate-templated lipid membranes to fluorescent solutes. Transmitted light and fluorescence images before and 10 minutes after the addition of (A) Alexa-647-labeled U15 RNA and (B) fluorescein isothiocyanate (FITC) for lipid-coated protamine sulfate coacervates (lipid composition: 37.9 mol.% DOPC, 30 mol.% DOPE, 30 mol.% DOPS, 2 mol.% DOPE-PEG 2kDa, 0.1 mol.% DOPE-Rh) in 1 mM MgCl_2_, 5 mM HEPES pH7.4 buffer.

No membrane imperfections were visible in the fluorescence rhodamine channels for the protamine sulfate coacervates to explain the differences between droplets that were permeable/impermeable to solute entry (Figure 4A-B), even when changing the focal plane to search the upper and lower portions of the structures. We interpret this as indicating any lipid membrane imperfections, where present, were below optical resolution. However, in the PDADMAC/PAA system (Figure 4 C-D), some microscale membrane defects were visible before addition of solutes and most of the droplets that accumulated solute had large regions without a membrane coating (Figure S3). Coacervates that accumulated FITC and RNA were also larger, indicating coalescence, which we attribute to the absence of an intact lipid membrane coating around those droplets. Differences between the two coacervate systems indicate a role for coacervate-specific lipid interactions in controlling the formation of the supported membranes, which is consistent with reported differences in solid supported lipid membranes on differently functionalized substrates.^15, 16, 59^ Cationic nanoparticles have been reported to result in pores/imperfections in supported lipid bilayer systems.^60^ The presence of pores/imperfections within the lipid membranes that surround coacervate droplets can be considered beneficial or detrimental, depending on the envisioned function. For example, while membrane barrier function relies on intact lipid bilayers, pores can facilitate solute uptake which could be useful for loading protocells that lack specific channels or transporters. Lipid-coated coacervate systems could be valuable in testing how polymer-lipid interactions generate membrane defects, which is important in cell biology and therapeutics.^61^

Transmitted light and fluorescence images before and 10 minutes after the addition of (C) Alexa-647-labeled U15 RNA and (D) fluorescein isothiocyanate (FITC) for lipid-coated PDADMAC/PAA coacervates (Lipid composition: 25 mol. % POPS, 50 mol. % POPC and 22.9 mol. % POPE, 2 mol.% DOPE-PEG 2kDa and 0.1 mol.% DOPE-Rh) in 100 mM NaCl, 10 mM Tris pH 7.4 buffer. Fluorescence signals are false-colored (DOPE-Rh: red, U15: blue, FITC: green) to aid in visualization. Note that different coacervates systems have different buffer and imaging conditions.

Another hallmark function of lipid vesicles and cell membranes is their ability to protect their interior components.^54^ In order to probe the ability of membranes formed around coacervates to protect their coacervate phase interior, the protease trypsin (MW 24 kDa) was added to the exterior solution following vesicle formation. Trypsin cleaves the peptide bonds of arginine and lysine residues,^62^ and as protamine sulfate is composed largely of arginine residues, multiple sites for cleavage exist. Trypsin cleavage of protamine sulfate, resulting in shorter peptide chains, led to coacervate dissolution within 3 minutes in the absence of lipid membranes (Figure 5A). Coacervate droplets encapsulated within GVs, however, were protected from trypsin cleavage and dissolution (Figure 5B).

**Figure 5:**
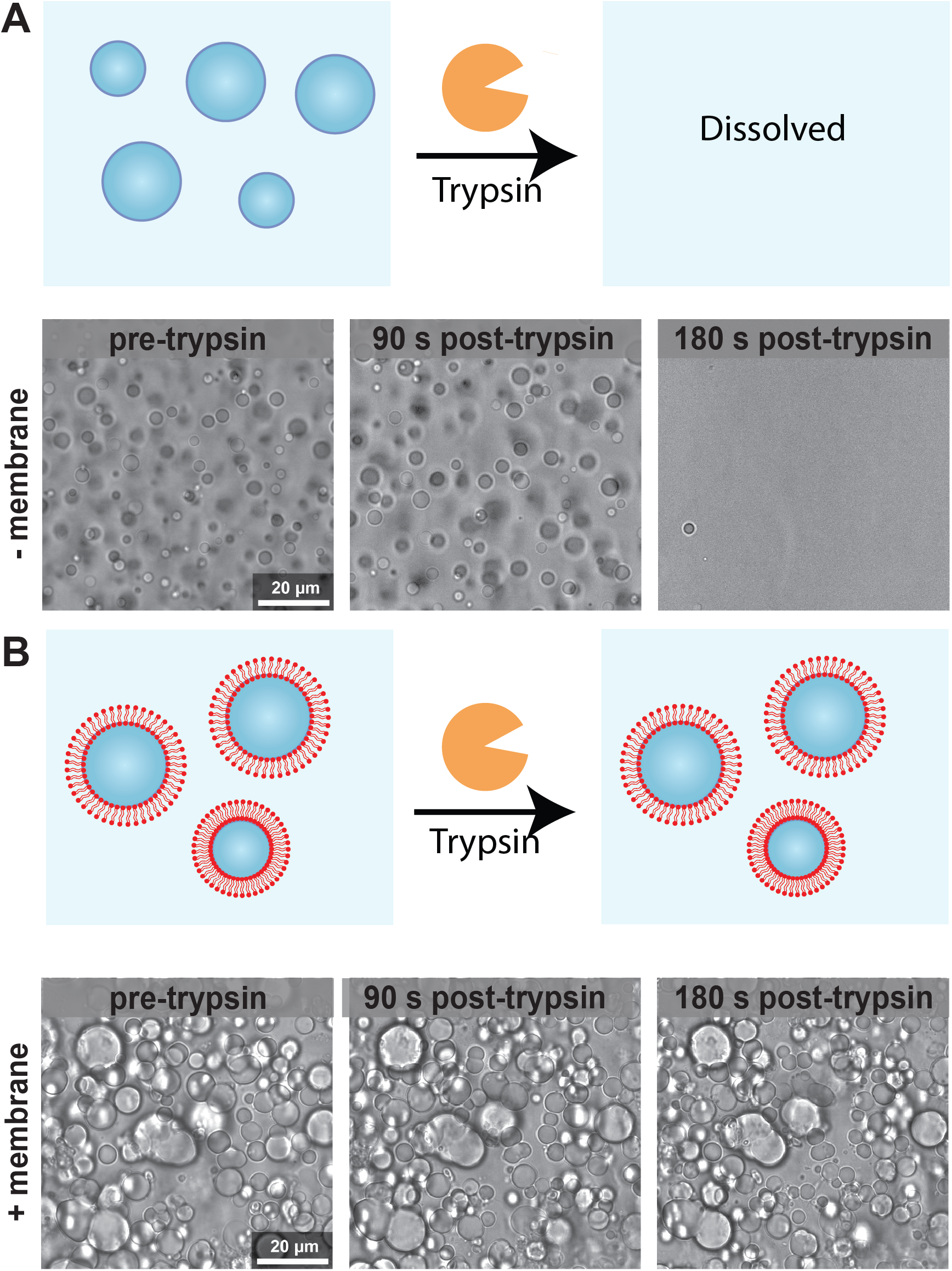
GVs membranes provide protection of interior coacervate phase from enzyme degradation. Transmitted light images of (A) 20 mM protamine sulfate coacervate droplets in solution and (B) GVs containing a protamine sulfate coacervate interior before and after addition of the protease trypsin. In the absence of protective lipid membranes, trypsin cleaves protamine sulfate triggering coacervate dissolution, while lipid membranes protect the protamine sulfate interior from trypsin degradation. Lipid composition of 37.9 mol% DOPC, 30 mol% DOPE, 30 mol% DOPS, 2 mol% DOPE-PEG 2kDa, 0.1 mol% DOPE-Rh was used for protamine in 1 mM MgCl_2_, 5 mM HEPES pH7.4 buffer.

### Encapsulation of biomolecules

Membrane-coated coacervate droplets are appealing both as models for biological cells, and as versatile artificial cells offering a dense, crowded cytoplasm surrounded by a lipid membrane. Macromolecule loading into cell-sized, “giant” lipid vesicles is challenging, with low and often heterogeneous encapsulation efficiency typical for traditional lipid hydration methods.^32, 63^ Microfluidic approaches offer excellent encapsulation and vesicle-to-vesicle homogeneity but require specialized methods and equipment.^64, 65^ Here, we evaluate macromolecule encapsulation in membrane-coated coacervates as a very simple alternative to loading by traditional vesicle hydration. We used 0.05 μM final concentration of U15 and BSA in the solution with and without PDADMAC/PAA coacervates (Figure 6). Traditional giant vesicles formed by gentle hydration exhibited low fluorescence intensities both inside and outside of the vesicles with an intensity ratio close to ~1 (Figure 6D and Figure S5). This indicates that the U15 and BSA were encapsulated at levels roughly equal to their external concentration (therefore estimated as equal to the added concentration of 0.05 μM). In contrast, membrane-coated coacervates formed by first incubating with the U15 or BSA, followed by membrane formation, showed strong accumulation of the labeled biomolecules. Intensity for labeled U15 was ~190 times (at least 9.4 ± 0.97 μM, a minimum value based on assuming external concentration is still 0.05 μM despite loss of solute to the coacervate phase) and BSA ~440 times (at least 22.1 ± 1.8 μM) that in the external solution (Figure 6A-D and Figure S6). Coacervate-templated vesicle formation provides a simple means of encapsulating biomolecules with excellent efficiency and high local concentrations, and with relatively low vesicle-to-vesicle variability as compared to other passive encapsulation methods.

**Figure 6:**
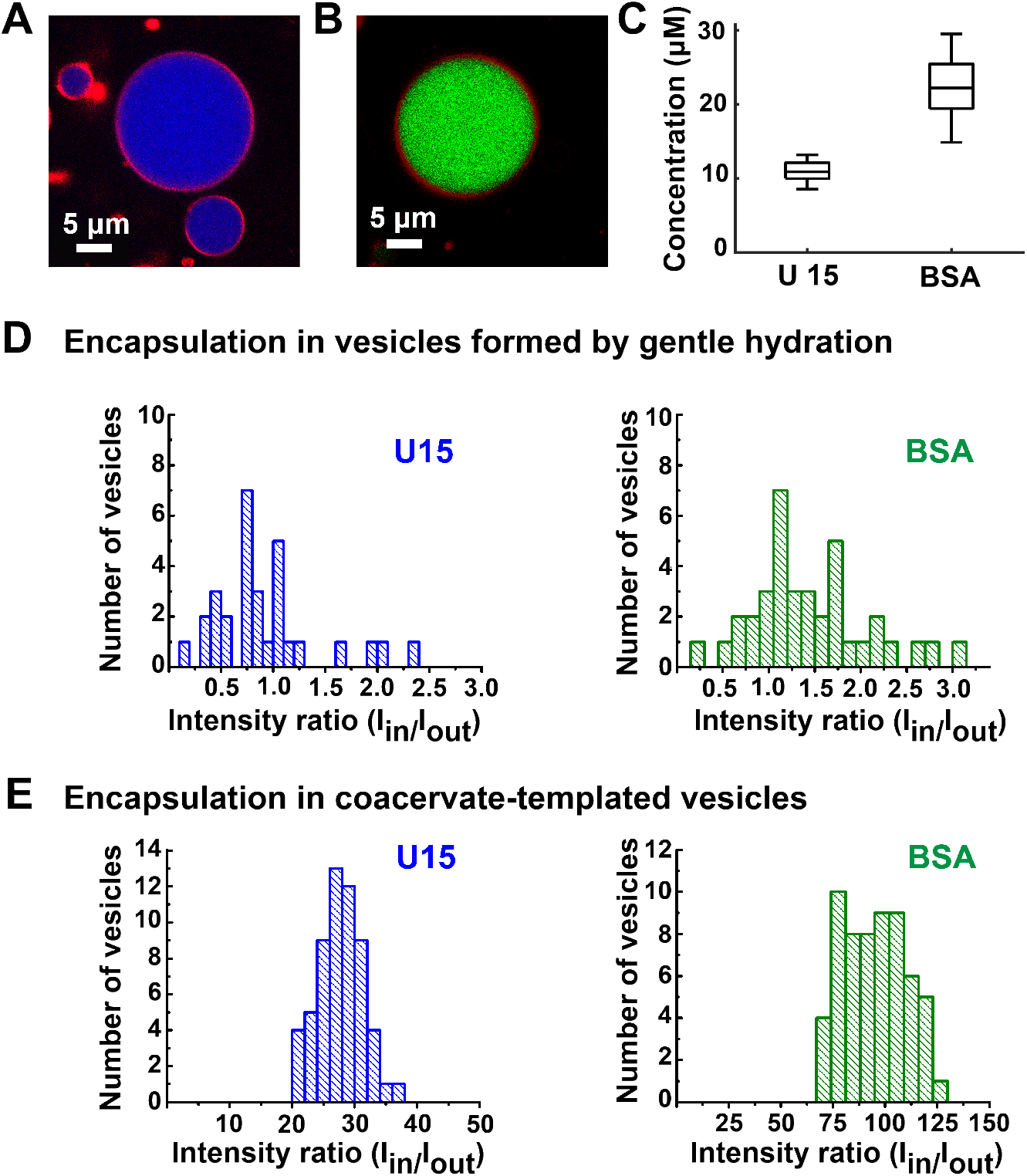
Encapsulation of biomolecules within coacervate templated vesicles. Representative images of coacervate-templated lipid vesicles encapsulating fluorescently-labeled oligonucleotide U15 RNA (A) or protein BSA (B). (C) Concentrations of U15 and BSA within the membrane templated coacervate droplets were plotted. Total solution concentration (moles added/total solution volume) for these experiments was 0.05 μM for both solutes. Center lines show the medians; box limits indicate the 25th and 75th percentiles; whiskers extend to the most extreme data points. Error bars show the standard deviation of multiple individual droplets obtained. (D, E) Intensity ratios for labeled U15 and BSA inside/outside the vesicles for traditional lipid vesicles formed by gentle hydration (D), and for coacervate-templated vesicles formed by hydration in the presence of coacervate droplets, (E). In both cases 0.05 μM of the U15 or BSA solute was present during membrane formation. The PDADMAC/PAA coacervate system in 100 mM NaCl, 10 mM Tris pH 7.4 buffer was used for these experiments. Lipid composition was 25 mol. % POPS, 50 mol. % POPC and 22.9 mol. % POPE, 2 mol.% DOPE-PEG 2kDa and 0.1 mol.% DOPE-Rh.

## Discussion

The work presented here shows that lipid membrane formation can be templated by complex coacervate droplets, forming what appear to be coacervate-supported lipid bilayers. Compared with giant lipid vesicle formation by gentle hydration in the absence of coacervates, the coacervate-templated membrane structures were more uniform in size, shape, and thickness (i.e. apparent lamellarity). Coacervate-supported membranes display rapid lipid diffusion, fluorescence polarization, and quenching behavior consistent with lipid bilayers. Within a population of lipid-coated droplets, some membranes appear to be complete, resisting uptake of solutes into the coacervate interior, while others within the same population are incomplete or imperfect, and allow solute uptake. Coacervate-templated membrane formation was observed for two different polyelectrolyte/lipid systems, indicating that this form of self-assembly is not limited to specific molecular interactions and likely relies largely on ion pairing interactions between the coacervate polycations and negatively charged lipid headgroups. Differences in the barrier function of membranes formed using different coacervate/lipid combinations point to the importance of chemical interactions between polyelectrolytes and lipids, most probably ion pairing between polycations and negatively charged lipid headgroups. It will be interesting to learn through future studies how these interactions can be tuned to design coacervate-supported lipid membranes for different levels of membrane perfection/permeability.

Coacervate-supported lipid membranes provide a new class of cell mimic that captures essential physicochemical features of both cell membranes and cytoplasm. No special equipment or techniques are necessary for production of these structures, and high loadings of biomacromolecules such as proteins or nucleic acids are readily achieved by allowing uptake into the coacervate droplets prior to membrane assembly. As such, these structures are appealing as artificial cells and could find use as model systems for studying macromolecule-membrane interactions-including those mediated by liquid-liquid phase separation-in the context of the macromolecularly-crowded cytoplasm of biological cells. Finally, this work points to the ease with which lipid membranes can form around coacervate droplets, potentially offering a straightforward route to accumulation and encapsulation of protobiomolecules to form protocells in prebiotic contexts. Membrane imperfections could be considered to have been beneficial, even necessary, for early compartments transitioning to protocells since effective transporter proteins would likely not yet have become available. Populations having both intact and imperfect membranes further offer opportunities for selective pressure to favor one or the other phenotype over time, depending on environmental stressors. Permeable membranes could also be beneficial in some artificial cell and biotechnology scenarios since they allow uptake without requiring special membrane protein transporters or pore-formers.

## Materials and Methods

### Materials

Protamine sulfate from salmon sperm was acquired from MP Biomedicals (Santa Ana, CA) or Sigma Aldrich. PDADMAC, PAA, HEPES, HEPES sodium salt, magnesium chloride hexahydrate (MgCl_2_.6H_2_O), trypsin from bovine pancreas, and fluorescein isothiocyanate (FITC) were purchased from Sigma Aldrich (St. Louis, MO). Lipids were acquired from Avanti Polar Lipids (Alabaster, AL): 1-palmitoyl-2-oleoyl-sn-glycero-3-phospho-L-serine (POPS), 1-palmitoyl-2-oleoyl-sn-glycero-3-phosphocholine (POPC), 1-palmitoyl-2-oleoyl-sn-glycero-3-phosphoethanolamine (POPE), DOPC (1,2-dioleoyl-sn-glycero-3-phosphatidylcholine), DOPE (1,2-dioleoyl-sn-glycero-3-phosphotidylethanolamine), DOPS (1,2-dioleoyl-sn-glycero-3-phosphotidylserine), DOPE-rhodamine, and DOPE-PEG MW 2 kDa. The fluorescently-labeled oligonucleotide, U15 RNA, was 5’-labeled with Alexa Fluor 647 (NHS ester) and purchased from Integrated DNA Technologies (Coralville, IA). Chemicals were used as received.

### Coacervate-supported membrane formation

#### Protamine sulfate coacervates

A 20 mM stock solution of protamine sulfate was prepared in 5 mM HEPES and 1 mM MgCl_2_ at pH 7.4. Giant vesicles were produced using the well-described gentle hydration method,^66, 67^ modified by using the 20 mM protamine sulfate. The lipid mixture was used for all protamine sulfate experiments: 37.9 mol.% DOPC, 30 mol.% DOPE, 30 mol.% DOPS, 2 mol.% DOPE-PEG 2kDa, 0.1 mol.% DOPE-Rh were dried within a glass tube to form a thin lipid film. 0.1 % DOPE-rhodamine was used for fluorescence microscopy. Then, the hydrating solution was added slowly down the sides of the test tube, and the solution was heated at 40 °C for 48 hours. For comparison between protamine sulfate containing samples versus giant vesicles, prepared giant vesicles used same lipid composition (37.9 mol.% DOPC, 30 mol.% DOPE, 30 mol.% DOPS, 2 mol.% DOPE-PEG 2kDa, 0.1 mol.% DOPE-Rh) and buffer conditions (5 mM HEPES and 1 mM MgCl_2_ at pH 7.4.). Note that we observed variability from batch-to-batch for protamine and the reason for that might be partly due to the sulfate not being a quality control priority in manufacture. However, sulfate is required for phase separation due to surface charge.

#### PDADMAC/PAA coacervates

PDADMAC/PAA at 1:1 ratio coacervate samples were prepared in charge concentration of 20 mM (charge of the molecule×concentration of the molecule = charge concentration) in HPLC grade water and 100 mM NaCl, 10 mM Tris at pH 7.4. Giant vesicles were produced using the well-described gentle hydration method,^66, 67^ modified by using the charge concentration of 20 mM PDADMAC/PAA solution as the hydrating phase. The lipid mixture was used for all PDADMAC/PAA experiments: 25 mol. % POPS, 50 mol. % POPC and 22.9 mol. % POPE, 2 mol.% DOPE-PEG 2kDa and 0.1 mol.% DOPE-Rh were dried within a glass tube to form a thin lipid film. 0.1 % DOPE-rhodamine was used for fluorescence microscopy. Then, the hydrating solution was added slowly down the sides of the test tube, and the solution was heated at 40 °C for 48 hours. For comparison between PDADMAC/PAA containing samples versus giant vesicles, prepared giant vesicles used same lipid composition (25 mol. % POPS, 50 mol. % POPC and 22.9 mol. % PE, 2 mol.% DOPE-PEG 2kDa and 0.1 mol.% DOPE-Rh) and buffer conditions (100 mM NaCl, 10 mM Tris at pH 7.4.).

#### Fluorescence Recovery After Photobleaching

Fluorescence recovery after photobleaching (FRAP) studies were conducted on vesicle membranes with an excitation at 543 nm for rhodamine-labeled lipids (DOPE-Rh). For each vesicle analyzed, a circular region of interest (ROI) (2μm) was chosen that contained a portion of the vesicle membrane without any aggregates or additional lipid bodies. 10 frames were collected pre-bleach, followed by 20 bleach frames at 100% laser power across all excitation wavelengths (458, 476, 488, 514, 543, and 633 nm), followed by 300 post-bleach frames. We used same procedure we reported^13^ and see Supplementary Information for details.

## Acknowledgment

This work was supported by the National Science Foundation, grants MCB-1715984 (protamine sulfate coacervate system) and NSF-RoL-RAISE 428-21 62G2 (PDADMA/PAA coacervate system).

## Author Contributions

F.P.C, A.M.M. and C.D.K designed the systems. F.P.C. performed coacervate-templated membrane formation (PDADMA/PAA and protamine sulfate systems), membrane quenching, permeability, encapsulation, and FRAP experiments. A.M.M. first observed the lipid assembly during hydration of lipid films and performed protamine sulfate coacervate-templated membrane formation, permeability, trypsin degradation protection, and FRAP experiments. C.D.K and F.P.C wrote the manuscript with contributions from A.M.M.

## Additional Information

The Supporting Information is available free of charge at. Materials and method, fluorescence images of lipids with PDADMAC/PAA, differential interference contrast and fluorescence images of protamine sulfate coacervates, fluorescence images of pore formation, fluorescence images of NBD-PE lipids, differential interference contrast and fluorescence images of encapsulation in GVs and coacervate templated GVs, yield calculation tables.

---

For Table of Contents Only:

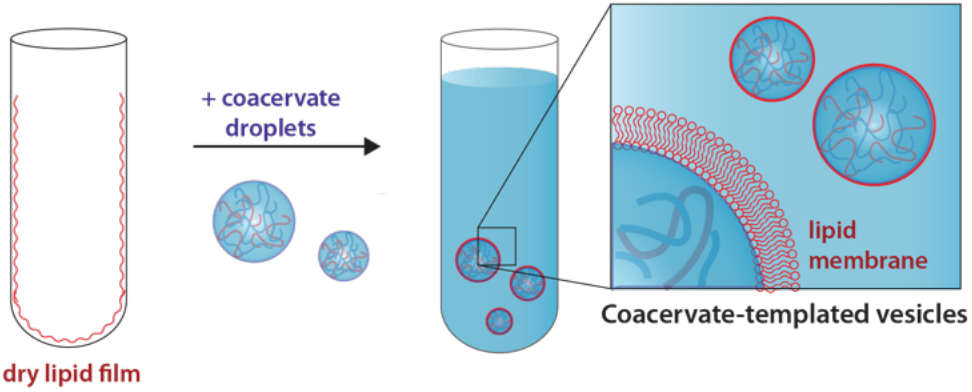

